# Transparency in Authors’ Contributions and Responsibilities to Promote Integrity in Scientific Publication

**DOI:** 10.1101/140228

**Authors:** Marcia McNutt, Monica Bradford, Jeffrey Drazen, Brooks Hanson, Bob Howard, Kathleen Hall Jamieson, Véronique Kiermer, Michael Magoulias, Emilie Marcus, Barbara Kline Pope, Randy Schekman, Sowmya Swaminathan, Peter Stang, Inder Verma

**Author notes:** Several authors’ employers are ORCID member organizations. Véronique Kiermer serves as Chair of the ORCID Board of Directors in a volunteer capacity. Contributions: Conceptualization, Supervision, Writing – original draft, Writing – review & editing. Contributions: Conceptualization, Writing – review & editing. Contribution: Conceptualization. Contributions: Conceptualization, Funding acquisition, Project administration, Resources, Writing – review & editing. Contributions: Conceptualization, Project administration, Resources, Writing – review & editing.

## Abstract

In keeping with the growing movement in scientific publishing toward transparency in data and methods, we argue that the names of authors accompanying journal articles should provide insight into who is responsible for which contributions, a process should exist to confirm that the list is complete, clearly articulated standards should establish whether and when the contributions of an individual justify authorship credit, and those involved in the generation of scientific knowledge should follow these best practices.

To accomplish these goals, we recommend that journals adopt common and transparent standards for authorship, outline responsibilities for corresponding authors, adopt the CRediT (Contributor Roles Taxonomy)^1^ methodology for attributing contributions, include this information in article metadata, and encourage authors to use the digital persistent identifier ORCID.^2^ Furthermore, we suggest that research institutions have regular open conversations on authorship criteria and ethics and that funding agencies adopt ORCID and accept CRediT. Scientific societies should further authorship transparency by promoting these recommendations through their meetings and publications programs.

## Challenges and Opportunities

The increase in international and interdisciplinary collaboration and the appearance of papers with hundreds of authors have complicated authorship practices in scientific publishing. At the same time, as pioneering journals have demonstrated, the digital revolution has made it possible to collect and quickly access author metadata. In response to these new challenges and opportunities, the science community should systematically address authorship practices that are too often obfuscated by uncertainty and confusion.

Publications are an important measure of research productivity for scientists across all disciplines. Publications, and citations to them, are in fact “the coin of the realm.” Despite the importance of scientific authorship to the professional prospects of scientists, the concept of “authorship” remains clouded. Does the order of authorship signify a descending level of contribution to the work? Not always. Is the first author responsible for verifying that all authors have acted responsibly? Not necessarily. Neither does the list of authors reveal to whom questions on specific aspects of the research should be directed nor who should be credited for a particularly groundbreaking aspect of the paper. Having an individual’s name listed among the authors does not even guarantee that the person contributed anything to the research. Nor does the specification of authors necessarily include all those who made substantive contributions to the work. The lack of openness and transparency on authorship stands in stark contrast to the rapid progress that journals have made on a number of other issues, such as access to data, materials, samples, and code.

These concerns prompted the National Academy of Sciences President Marcia McNutt to convene a small group from leading journals and scientific organizations at the Annenberg Retreat at Sunnylands in Rancho Mirage, California, in February 2017 to discuss how to promote standards that would increase transparency in author contributions to research papers in the natural and social sciences. Because these organizations represented serve a global scientific audience, we believe the proposed recommendations could apply internationally.

The retreat was premised on the assumptions that a more transparent and standardized approach to authorship might prove useful to (1) authors in clarifying their contributions and roles; (2) proposal evaluation, hiring, tenure and promotion, and awards committees in assigning credit to authors for their contributions in multi-authored papers; (3) investigators in determining culpability in cases of scientific misconduct; (4) readers in directing queries on data, materials, methods, and other questions about the research; (5) researchers and program managers in preserving more granular data on how complex science is conducted; and (6) research organizations, including universities and centers, in establishing authorship protocols for their scholars.

## Promoting Practices That Protect the Integrity of Authorship

The notion of authorship implies both credit and accountability.^3^ Yet the authorship conventions vary across disciplines,^4, 5, 6^ across cultures internationally,^7^ and even between research groups/labs. The various conventions include expectations for what effort earns authorship, what the order of authorship signifies (if anything), how much accountability for the research the corresponding author assumes, and the extent to which authors are accountable for aspects of the work that they did not personally conduct. Authorship practices that are considered detrimental to research^8^ are

- “ghost authors”^9^ (authors who contributed to the work but are not listed, generally in order to hide a conflict of interest from editors, reviewers, and readers);
- “guest/gift/honorific authors”^10^ (individuals given authorship credit who have not contributed in any substantive way to the research but are added to the author list by virtue of their stature in the organization);
- “orphan authors” (authors who contributed materially to the work but are omitted from the author list unfairly by the drafting team); and
- “conscripted authors” (unwitting authors who had no part in the work but whose names are appended to the paper without their knowledge to increase the likelihood of publication).

To protect the integrity of authorship, a number of journals have published author disclosure policies.^11^ These statements generally require those identified as authors to specify their contributions to the final work, sometimes using pre-set categories. The resulting self-declarations may be published as a statement within the article, for example, in the acknowledgments. However the form of these contribution statements and the ways in which they are presented/embedded in a paper vary widely. Because the disclosures are at present rarely captured in the journal metadata, the record of author contribution cannot readily be independently confirmed and aggregated on curricula vitae or as part of the author’s ORCID record. At the same time, the common practice of linking the identity of the author to the contribution by initials only makes it impossible to search the record electronically. Capture via the metadata could expedite preparation of proposals and employment applications and provide more granular information for tenure and promotion committees. Such standardized metadata would also facilitate the study of authorship practices and an evidence-based approach to their influence on research assessment.

A more transparent system of declaring author contribution will protect the integrity of science by

- expediting reader requests for access to samples, code, data, materials, or more information on experimental methods;
- limiting collateral damage if certain restricted elements of the study are found to be unreliable;
- ensuring fairness in that all authors receive credit for their actual contributions; and
- discouraging or preventing unethical authorship practices.

We therefore make the following recommendations for achieving a more transparent system of assigning author credit for contributions in the scholarly literature.

## Recommendations for Journals

The following recommendations are best practices currently in use in leading scientific journals.

### Set standards for authorship

Journals should increase authorship transparency by promulgating standards specifying the contributions that warrant authorship, indicating the responsibilities that are entrusted to corresponding authors (CAs), and implementing cross-publisher standardized electronic methods of capturing author contributions in journal metadata. As the first step, we recommend that journals adopt the following statement as a best disclosure practice for crediting all authors of a paper:

> Each author is expected to have made substantial contributions to the conception or design of the work; or the acquisition, analysis, or interpretation of data; or the creation of new software used in the work; or have drafted the work or substantively revised it; AND has approved the submitted version (and any substantially modified version that involves the author’s contribution to the study); AND agrees to be personally accountable for the author’s own contributions and for ensuring that questions related to the accuracy or integrity of any part of the work, even ones in which the author was not personally involved, are appropriately investigated, resolved, and documented in the literature.

This statement is adapted from a similar one developed by the International Committee of Medical Journal Editors (ICMJE) that is widely used within the medical publishing community. The statement has been generalized to encourage broader adoption.

### Provide expectations for corresponding authors

Second, journals should clearly articulate their expectations for CAs. The CA should ensure that all listed authors have approved the manuscript before submission to a journal and that all authors receive the submission and all substantive correspondence with editors, as well as the full reviews. Journals should ask the CAs to verify that all data, materials (including reagents), and code, even those developed or provided by other authors, comply with the transparency and reproducibility standards of both the field and journal. This responsibility includes but is not limited to (1) ensuring that original data/materials/code upon which the submission is based are preserved and retrievable for reanalysis; (2) confirming that data/materials/ code presentation accurately reflects the original; and (3) foreseeing and minimizing obstacles to the sharing of data/materials/code described in the work. In short, the CA should be responsible for managing these requirements across the author group and ensuring that the entire author group is fully aware of and in compliance with best practices.

Journals might also task the CAs with fulfilling other requirements for publication (e.g., verifying that the appropriate research oversight review committees have approved the research and confirming that any required Material Transfer Agreements were obtained). It is likely that the CAs will be responsible for delivering the necessary publication paperwork from coauthors (e.g., competing interest forms, author contribution forms, funding information, etc.).

To discourage the unethical authorship practices described above (see Promoting Practices That Protect the Integrity of Authorship), journals should also require that CAs reveal as appropriate whether the manuscript benefited from the use of editorial services that, if undeclared, might constitute an undisclosed conflict of interest. Examples include use of an editor from an organization that may have a vested interest in slanting the results or reliance on a technical writer at a level that would warrant authorship credit. These situations might variously be addressed by including a statement in the acknowledgments, by describing the effort in the methods section, or by adding an author. Many journals require CAs to indicate whether any authors on earlier versions have been removed or new authors added and why. This simple step discourages the practice of guest authors or orphan authors. CAs should also assume responsibility for ensuring that all authors who deserve to be credited on the manuscript are indeed identified, that no authors are listed who do not deserve authorship credit, and that author contributions, where they are provided, are expressed accurately. It is incumbent on the corresponding author to ensure that all authors (or group/lab leads in large collaborations) have approved the author list and contribution description.

### Commit to the use of CRediT

Our third recommendation for journals is that they commit to use or, where appropriate, adapt CRediT^12^ (Contributor Role Taxonomy) as, in our estimation, the best available method for embedding authors’ contributions in journal metadata. Because CRediT was designed by a stakeholder group consisting of practitioners and researchers from the physical, life, and social sciences, it captures broadly the perspectives of research publishing in these domains. The 14 CRediT categories for contributor roles are evidence based in that they were distilled from free-form author statements and acknowledgments. Insofar as CRediT is experiencing increasing uptake by the author community, adopting or adapting this standard is a better approach than devising a new one.

Uniformity of declarations across journals is in the authors’ best interests because such statements will not need to change if the same paper is rejected and submitted to another journal or if the same research team with similar roles submits a follow-on study to the same or another outlet. Journals’ adoption of CRediT in both human- and machine-readable forms, that is as part of the publication and its metadata, will facilitate transparency of author contributions in different contexts—via syndication, indexing services, and possibly future applications.

Moreover, asking the authors to agree on author contribution statements as the manuscript is finalized is likely to be a more accurate record of what transpired than asking participants to recall the information post facto (e.g., when authors are considered for promotion, research grants, or major prizes). CRediT offers a standard vocabulary and a consistent framework to facilitate authorship discussions amongst contributors to a study. In this regard, it is important to note that the taxonomy includes but is not limited to traditional authorship roles. It is intended to describe authors’ contributions within the framework of authorship standards, not to define such standards. Adopting CRediT would also help alleviate some of the confusion across disciplines and cultures regarding meaning in author order. These differences are currently so ingrained that forcing a common standard is impractical and will be unnecessary if CRediT becomes widely adopted.

We recommend that the CRediT roles be embedded within author metadata rather than as a separate paragraph of text, often linked just to author initials. This would allow contribution information to be transmitted to abstracting and indexing services and other databases such as CrossRef and ORCID and provide a common machine-readable format across journals. We recognize that full implementation will require some further development of common schemas and standards across journals, organizations, and communities.

We recognize that adoption of CRediT by journals is an aspirational goal but believe that now is the time, in the early phases of journal experimentation with attributing contributions to authors, to champion this standard and its widespread adoption. We note that there are significant benefits in some of the formats currently implemented (e.g., note the specificity in author contributions in a recent *Nature* article^13^). Use of CRediT need not be the only approach for a journal to capture how each author earned the right to be listed (e.g., additional information can be captured in acknowledgments or footnotes), but if CRediT is available consistently, in machine-readable form and through the journal metadata, author contributions will transition from hearsay to quantifiable evidence.

### Encourage authors to adopt ORCID and other standard identifiers

Finally, we believe that all journals in the physical, life, and social sciences should strongly encourage author adoption of ORCID. In order to eliminate name confusion and ensure appropriate attribution of publications and citations to the correct authors, many major journals are already requiring ORCID iDs for first, corresponding, or all authors. When combined with CRediT, the potential exists to reliably link author records to publications, to capture author contributions in the journal’s metadata, and to track and retrieve an individual’s authorship contributions across publications and across time. Although it cannot alone guarantee secure identity, adoption of ORCID is one more check against author identity fraud.

Other persistent identifiers (PIDs) have emerged to uniquely identify different funders of research, and a multi-stakeholder collaboration is ongoing to establish standard PIDs for institutions. At the same time, some fields have adopted PIDs for samples. Development is under way to explore or extend PIDs for repositories and instruments that would lead to further transparency and integrity. We encourage journals to follow these developments and prepare for wider adoption when standards emerge.

## Recommendations for Other Stakeholders

The focus of the Sunnylands retreat was on the role of journals because most of those in attendance represented that sector. However, editors and publishers realize that they are but one group involved in the process of protecting the integrity of authorship.

### Universities/research institutions

Universities and other research institutions play a central role as well. Recognizing that these institutions house most of those whose scholarly work appears in journals, we recommend that departments and other research organizations develop, post, distribute, and regularly review and update their policies on authorship. This process should actively involve faculty/investigators, postdoctoral scholars, graduate and undergraduate students, staff, and any others making important contributions to scholarship in the sciences. Questions answered by these policies would include how the organization or a research team within it decides which contributions warrant first authorship, co-authorship, acknowledgments, or no mention.

We recognize that these decisions vary with the culture of various disciplines and nations. But transparency in how the decision is made before the research is undertaken can avoid later conflicts that journals are ill equipped to mediate.

Universities/research institutions worldwide should encourage open conversations, at least annually, at the departmental level about both the policy and criteria for authorship. The goals of this recommendation include familiarizing students, postdoctoral scholars, and new employees (including foreign visitors) with the norms of the field; standardizing criteria among and across laboratories within the same department; and making all parties aware of norms characterizing the production and crediting of science. To ensure that the same criteria for authorship are applied to personnel from different laboratories, universities should expect principal investigators involved in multi-laboratory collaborations (even if across universities) to engage in the same discussion. When two students make the same level and kind of contribution to a collaborative study, their status as author should not depend on the lab with which the student is affiliated.

### Funding agencies

Funding agencies could easily tip the scales toward widespread acceptance of a truly international standard for the unique identity of scholars by adopting ORCID as the preferred/default system for their principal investigators. If every investigator supported by U.S. and international funding agencies was required to use an ORCID iD, the standard would become the norm. A positive step in this direction was the recent decision by the European Commission to recommend the adoption of ORCID as its preferred system of unique identifiers for its 9th Framework Programme.^14^ Additionally, it would be helpful if in requests for bibliographical references in support of an application as well as in progress reports, funding agencies were to accept representation of the applicant’s contributions to the work as expressed via CRediT—for example, by accepting citation formats that allow the expression of CRediT terms so as to capture the applicant’s contribution to collaborative research. In the near future, support from funding agencies will be required to underwrite efforts to seamlessly integrate CRediT, ORCID, journal metadata, and scholarly archives, including public archives.

### Scientific societies

Many scientific societies have strongly endorsed efforts in transparency through editorials in their journals, special sessions at their meetings, and policies in their own publishing programs. We suggest that they build on these efforts by organizing special sessions on the topic of integrity in authorship at scientific meetings to discuss issues relevant to each society. Each organization should consider how the society can move briskly in its own journal publishing program to implement the recommendations in this white paper. These should include updates to their author instructions and guidelines.

## Further Steps

We recognize that the path we have outlined, while a leap forward in authorship transparency, leaves a number of vexing issues unresolved. CRediT provides a taxonomy of contributor roles, but does not allow for a hierarchy much less quantification of contributions by those on a research team: Who took the lead on the data analysis? Who provided the most significant insight in the data interpretation? Furthermore, the broad categories of contribution do not allow for additional detail on exactly what each author did. If three authors provided modeling, which one created the ocean model, which one conceived the atmospheric model, and which one provided the carbon model?

To address such questions, we encourage the creation of a cost-free, Web-based tool that facilitates author experimentation about ways to increase transparency and accountability in authorship beyond what is currently possible with CRediT. This tool would need to build authorship lists and contribution statements containing a certain minimum threshold of information required by journals. We propose this information would include ORCID iDs and the machine- and human-readable CRediT roles. The tool could also allow authors to experiment by

- assigning microcredits (e.g., attributing tables, figures, etc. to individual authors);
- establishing leading and supporting roles in each of the CRediT categories;
- adding further narratives to provide more context for the contribution (e.g., exactly which data collection, which code development, etc.);
- allowing authors to put percentages of effort in each category (e.g., “I spent 25% of my time on the project in each of the following categories: data collection, data analysis, data interpretation, and writing.”); and
- allowing the team to assign percentages of effort to each of the contributors in each category (e.g., “Smith did 90% of the project design, and Jones did 10% of the project design.”).

Authors could experiment with this tool and gain experience in attributing beyond the simple implementation of CRediT—for example, by annotating the contributions to match a specific figure. As long as the tool exports the minimal information required by a journal, in a format that the journal can receive, it could still be useful in the traditional workflow. Journals may decide to preserve this additional information in acknowledgments, footnotes, or by other means.

We also recognize that additional development is needed for export of CRediT to ORCID records. This step should be undertaken as soon as practical.

## Conclusions

Research publications are not only a primary vehicle through which scientists communicate knowledge but also a way in which the scholarly community establishes who deserves the credit or, in rare cases, the blame for the reported work. Presently the connection between authorship and a scientist’s contributions is poorly captured for posterity, nonstandard at best, and lacking integrity at worst. The recommendations we have made to increase the integrity of authorship are practical, able to be implemented in short order, and provide a path toward increasing transparency.

## Acknowledgments

The authors are grateful to The Annenberg Foundation Trust at Sunnylands (David Lane, President) for sponsoring this retreat.

